# Cardiovascular and Locomotor Effects of Binary Mixtures of Common “Bath Salts” Constituents: Studies with Methylone, MDPV, and Caffeine in Rats

**DOI:** 10.1101/2024.01.31.578069

**Authors:** Robert W. Seaman, David G. Galindo, Benjamin T. Stinson, Agnieszka Sulima, Kenner C. Rice, Martin A. Javors, Brett C. Ginsburg, Gregory T. Collins

## Abstract

**Background and Purpose:** The use of “Bath Salts” drug preparations has been associated with high rates of toxicity and death. Preparations often contain mixtures of drugs including multiple synthetic cathinones or synthetic cathinones and caffeine; however, little is known about whether interactions among “Bath Salts” constituents contribute to the adverse effects often reported in users.

**Experimental Approach:** This study used adult male Sprague-Dawley rats to characterize the cardiovascular effects, locomotor effects, and pharmacokinetics of methylone, MDPV, and caffeine, administered alone and as binary mixtures. Dose-addition analyses were used to determine the effect levels predicted for a strictly additive interaction for each dose pair.

**Key Results:** Methylone, MDPV, and caffeine increased heart rate and locomotion, with methylone producing the largest increase in heart rate, MDPV producing the largest increase in locomotor activity, and caffeine being the least effective in stimulating heart rate and locomotor activity. MDPV and caffeine increased mean arterial pressure, with caffeine being more effective than MDPV. The nature of the interactions between methylone and MDPV tended toward sub-additivity for all endpoints, whereas interactions between MDPV or methylone and caffeine tended to be additive or sub-additive for cardiovascular endpoints, and additive or supra-additive for increases in locomotion. No pharmacokinetic interactions were observed between individual constituents, but methylone displayed non-linear pharmacokinetics at the largest dose evaluated.

**Conclusion and Implications:** These findings demonstrate that the composition of “Bath Salts” preparations can impact both cardiovascular and locomotor effects and suggest that such interactions among constituent drugs could contribute to the “Bath Salts” toxidrome reported by human users.

**What is already known:** “Bath Salts” preparations are associated with a sympathomimetic toxidrome in human users.

**What this study adds:** Characterization of both pharmacokinetic and pharmacodynamic interactions between common “Bath Salts” constituents with regard to cardiovascular and locomotor effects.

**Clinical Significance:** The vast majority of drug overdose deaths involve more than one substance. Though these studies focused on combinations of stimulant drugs, they provide direct evidence that the toxidrome resulting from multi-drug overdoses can be significantly different than would be expected for a single drug.

## INTRODUCTION

Synthetic cathinones became popular in the early 2000s in the form of “Bath Salts” preparations due to their fleeting status as legal alternatives to prototypical illicit stimulants such as cocaine and methamphetamine. They are associated with several bizarre and adverse effects as well as a relatively large number of emergency room visits. Individuals reporting “Bath Salts” use exhibited both psychiatric (paranoia, hallucinations, agitation) and physiological (tachycardia, hyperthermia) symptoms that proved to be lethal in some cases (Kyle et al., 2011; Spiller et al., 2011; Murray et al., 2012; Johnson and Johnson, 2014). Chemical analyses of the initial wave of “Bath Salts” preparations suggested that most preparations contained 3,4-methylenedioxypyrovalerone (MDPV; a selective dopamine and norepinephrine transporter reuptake inhibitor), 3,4-methylenedioxymethcathinone (methylone; a non-selective monoamine transporter substrate), or 4-methylmethcathinone (mephedrone; a non-selective monoamine transporter substrate) (Spiller et al., 2011; Shanks et al., 2012; Seely et al., 2013). Consistent with their popularity as recreational drugs, the discriminative stimulus effects of MDPV and methylone overlap with those of cocaine and methamphetamine (Fantegrossi et al., 2013; Gatch et al., 2013; Collins et al., 2016; DeLarge et al., 2017; Harvey et al., 2017; Smith et al., 2017; Gatch and Forster, 2020; Seaman et al., 2021), and both drugs are readily self-administered by rats (Aarde et al., 2013; Watterson et al., 2014; Schindler et al., 2016a; Gannon et al., 2017, 2018c) and non-human primates (Collins et al., 2019; de Moura et al., 2021). Since their emergence, the United States Drug Enforcement Agency has placed 13 synthetic cathinones, including MDPV, methylone, and mephedrone, under Schedule I regulations (Drug Enforcement Administration, Department of Justice 2011, 2014). Subsequently, clandestine laboratories have attempted to skirt these regulations by making slight chemical modifications to these structures, resulting in a large number of “legal” synthetic cathinone derivatives that retain activity as substrates for monoamine transporters and monoamine reuptake inhibitors. Although the general mechanisms of action remain intact, the introduction of modest structural differences results in differences in the potencies and selectivities of these compounds to interact with monoamine transporters, resulting in changes in abuse-related effects, including discriminative stimulus and reinforcing effects (Gannon et al., 2018c, 2018a; Seaman et al., 2021).

Though the constantly evolving landscape of novel synthetic cathinones entering the drug market likely contributes to the heterogeneity of “Bath Salts” preparations across time, one constant is that “Bath Salts” preparations are most often mixtures of multiple synthetic cathinones (e.g., methylone and MDPV), or mixtures of synthetic cathinone(s) and other psychoactive substances (e.g., caffeine). The inclusion of caffeine in “Bath Salts” preparations is likely influenced by the fact that it is legal, relatively inexpensive, and capable of producing mild stimulant properties that are mediated by interactions between adenosine and dopamine systems (Ferré, 2016). Though the degree to which the subjective and/or adverse effects of “Bath Salts” preparations are influenced by caffeine is unclear, preclinical studies have begun examining the role of caffeine in the abuse-related effects of caffeine-containing drug mixtures. Indeed, drug discrimination studies in rats have demonstrated that the discriminative stimulus effects of caffeine not only overlap with those of cocaine and MDPV, but mixtures of cocaine+caffeine and MDPV+caffeine exhibit supra-additive interactions with respect to their cocaine-like discriminative stimulus effects (Collins et al., 2016; Seaman et al., 2021). Furthermore, interactions between MDPV and caffeine, and methylone and caffeine, have been demonstrated to be greater than additive with regard to reinforcing effectiveness, determined using progressive ratio and economic demand procedures in rats (Gannon et al., 2018b, 2019) as well as interact synergistically to reinstate extinguished responding for drug-paired stimuli (Doyle et al., 2021).

Although those studies highlight the potential for interactions amongst the abuse-related effects of common “Bath Salts” constituents, the extent to which this synergism extends to toxic effects remains unknown. Indeed, the physiological effects of MDPV in rats are consistent with prototypical stimulants (e.g., cocaine, methamphetamine), including dose-dependent increases in heart rate, blood pressure, and locomotion (Baumann et al., 2013; Schindler et al., 2016b; McClenahan et al., 2019; Concheiro et al., 2022). The effects of MDPV on body temperature are mixed, with some reports of mild hyperthermia and some studies reporting no effects on body temperature (Kiyatkin et al., 2015; Gannon et al., 2016, 2018d; Horsley et al., 2018). The physiological effects of methylone are less well described, however, it has been shown that methylone dose-dependently increases locomotor activity and produces a small hyperthermic effect in rodent models (López-Arnau et al., 2013; Elmore et al., 2017; Štefková et al., 2017; Javadi-Paydar et al., 2018; Centazzo et al., 2021).

There is some evidence suggesting that there could be synergism between the physiological effects of methylone and MDPV or between either cathinone and caffeine as has been seen with other stimulants (Derlet et al., 1992; Mehta et al., 2004; McNamara et al., 2007; Vanattou-Saïfoudine et al., 2010b, 2010a). For instance, both methylone and MDPV have been shown to be metabolized by cytochrome p450 2D6 (CYP2D6) in humans, and methylone has been suggested to be a mechanism-based inactivator of its primary metabolizing enzyme (Pedersen et al., 2013; Elmore et al., 2017), potentially culminating in a pharmacokinetic interaction between these two synthetic cathinones that could enhance the toxidrome produced by either drug alone. Although a pharmacokinetic interaction between either synthetic cathinone and caffeine would not be expected given the breadth of enzymes responsible for metabolizing caffeine that are largely non-overlapping with either cathinone (Kot and Daniel, 2008), the ability of caffeine to disinhibit dopamine D_2_ signaling through disruption of dopamine D_2_ and adenosine A_2A_ receptor heteromers (Ferré et al., 1997; Ferré, 2016) could result in a pharmacodynamic interaction between methylone or MDPV and caffeine to drive enhancements in adverse effects relative to what would be observed for either drug alone. It is also worth mentioning that we have previously reported an increased incidence of drug-related fatalities in rats that were allowed to self-administer binary “Bath Salts” mixtures containing methylone (i.e., methylone + MDPV and methylone + caffeine) (Gannon et al., 2018b, 2019), suggesting that these common “Bath Salts” constituents could interact to result in a more toxic and/or lethal drug preparation relative to methylone alone.

Whereas interactions between the reinforcing effects of binary mixtures of methylone, MDPV, and caffeine have been described, interactions between their physiological effects remain unknown. To this end, the current studies utilized radiotelemetry to characterize the physiological and pharmacodynamic effects of drugs alone and in combination, serial blood sampling and high-performance liquid chromatography with tandem mass spectrometry to characterize pharmacokinetic profiles of drugs alone, and in combination, and quantitative dose addition analytical approaches to define the nature of the interaction for heart rate, mean arterial pressure, and locomotion to test the following hypotheses: 1) methylone, MDPV, and caffeine will produce dose-dependent increases in heart rate, mean arterial pressure, and locomotor activity; 2) binary mixtures of methylone, MDPV, and caffeine will produce supra-additive increases in heart rate, blood pressure, and locomotor activity; 3) large doses of methylone will result in greater-than-expected blood levels of methylone, consistent with methylone displaying non-linear pharmacokinetics and acting as a mechanism-based inactivator of its metabolizing enzyme; and 4) greater than expected levels of MDPV will be observed when methylone and MDPV are co-administered, whereas blood levels of caffeine will be unaffected by co-administration of methylone.

## METHODS

### Subjects

120 male Sprague-Dawley rats (275–300 g upon arrival) were purchased from Envigo (Indianapolis, IN, United States) and maintained in a temperature- and humidity-controlled room. Rats were individually housed and maintained on a 14/10 h light/dark cycle. All experiments were conducted during the light cycle with sessions conducted at approximately the same time each morning. Rats were provided ad libitum access to Purina rat chow and water outside of experimental sessions. All studies were carried out in accordance with the Institutional Animal Care and Use Committees of the University of Texas Health Science Center at San Antonio and the eighth edition of the Guide for Care and Use of Laboratory Animals (National Research Council (United States) Committee for the Update of the Guide for the Care and Use of Laboratory Animals 2011).

### Surgeries

Rats were anesthetized with 2–3% isoflurane and prepared with either a single chronic indwelling catheter in the left femoral vein (for radiotelemetry studies; n=106), or two chronic indwelling catheters, one each in the left and right femoral veins (for the pharmacokinetic studies; n=14), as previously described (Seaman and Collins, 2021; Seaman et al., 2022). Catheters were tunneled under the skin and attached to a vascular access button placed in the mid-scapular region. Immediately following surgery, rats were administered Excede (20 mg/kg; SC) to prevent infection. Rats were allowed 5–7 days to recover during which time catheters were flushed daily with 0.5 ml of heparinized saline (100 U/ml). Thereafter, in rats being used for telemetry studies, incisions were made above the right femoral artery and on the abdomen to allow for a telemetric probe (model HD-S10; Data Sciences International, St Paul, MN) capable of collecting mean arterial pressure (MAP), heart rate (HR), core body temperature, and locomotor activity to be implanted in the abdominal cavity. Upon the placement of the probe, the pressure transducer was passed through the muscle and inserted into the femoral artery as described previously (Collins et al., 2011). Rats were allowed 5–7 days to recover from surgery prior to the beginning of testing.

### Drugs

3,4-methylenedioxypyrovalerone (MDPV) and 3,4-methylenedioxy-N-methylcathinone (methylone) were synthesized as racemic HCl salts by Agnieszka Sulima and Kenner Rice at the Intramural Research Program of the National Institute on Drug Abuse (Bethesda, MD, USA). Caffeine was purchased from Sigma-Aldrich (St. Louis, MO, USA). All drugs were dissolved in physiological saline. Drugs were administered via intravenous (IV) infusion in a volume of 1 ml/kg with the exception of doses of caffeine larger than 10 mg/kg which were administered at 2.3 ml/kg due to the solubility limit of caffeine in saline (∼14 mg/ml).

### Telemetry

#### Apparatus

Telemetry experiments were conducted in custom-built arenas with black walls and floors (43 cm x 56 cm x 38 cm) that were placed on receivers (Data Sciences International, St. Paul, MN) to allow for the collection of HR, MAP, core body temperature, and locomotor activity measures using Dataquest A.R.T. software.

### Drugs alone

#### Procedure

On test days, telemetry probes were switched on and rats were placed within testing arenas for one hour prior to IV administration of bolus doses of MDPV (0.032-3.2 mg/kg; n=10), methylone (0.32-10 mg/kg; n=9), or caffeine (0.32-32 mg/kg; n=11), followed by a 0.5 ml saline flush. Saline was evaluated first in all rats, after which all doses of a single drug were evaluated in a pseudorandomized order. Recording continued for three hours after administration of drug (or saline). Test sessions were conducted twice per week.

### Data Analysis

Heart rate, mean arterial pressure, and locomotor activity are presented as mean ± standard error of the mean (S.E.M.). Time courses were averaged into 10-min bins and analyzed via two-way repeated measures analysis of variance (ANOVA) with post-hoc Holm-Sidak tests comparing the effects of each dose to those of saline. Dose-effect curves are presented as the mean ±S.E.M. area under the curve (AUC) and were analyzed by one-way repeated-measures ANOVA with post-hoc Holm-Sidak tests.

### Drug Mixtures Procedure

On test days, telemetry probes were switched on and rats were placed within the testing arenas for one hour and subsequently intravenously infused with one dose-pair of mixtures of either methylone+MDPV (in ratios of 3:1 [n=9], 1:1 [n=8], or 1:3 [n=7]), MDPV+caffeine (in ratios of 3:1 [n=8], 1:1 [n=8], or 1:3 [n=9]), or methylone+caffeine (in ratios of 3:1 [n=8], 1:1 [n=9], or 1:3 [n=10]). Dose-pairs are listed in Table 1.

**Table 1:**
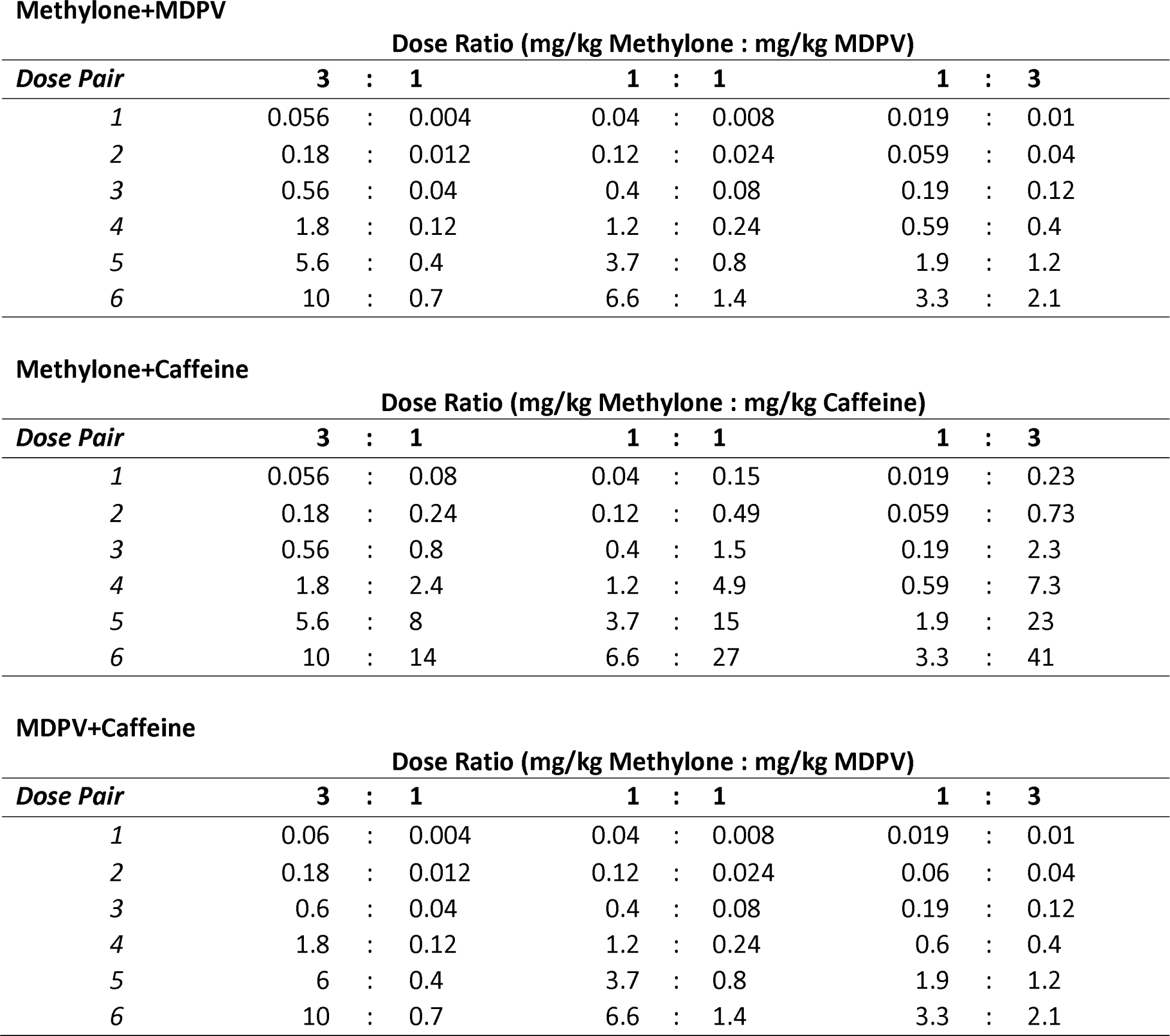
Dose pairs for each ratio of each binary mixture evaluated.

Recording continued for 3-h following drug administration. Rats only ever received one dose ratio of one drug mixture, and the order of dose-pairs evaluated was pseudorandomized, but the largest dose-pairs were always evaluated last. Drug infusions were immediately followed by an infusion of 0.5 ml of saline. Recording continued for three hours thereafter. Test sessions were conducted twice per week and saline was evaluated first in all rats.

### Data Analyses

Dose-effect curves for heart rate, blood pressure, and locomotor activity for individual drugs were fit using linear regressions of the data spanning the 20–80% effect levels, from which the ED_50_, slope, and y-intercept were obtained for individual subjects. Predicted effect levels for each dose-pair evaluated (listed in Table 2) were calculated based on the concepts of dose equivalence and dose addition (for review, see (Doyle et al., 2022)). Briefly, for a given endpoint (e.g., heart rate) and a given binary mixture (e.g., 1:1 MDPV+caffeine), the dose of the drug producing the lesser effect (DoseB) was converted to an equivalent dose of the drug producing the greater effect (DoseB_eq_A) using the following equation:

**Table 2:**
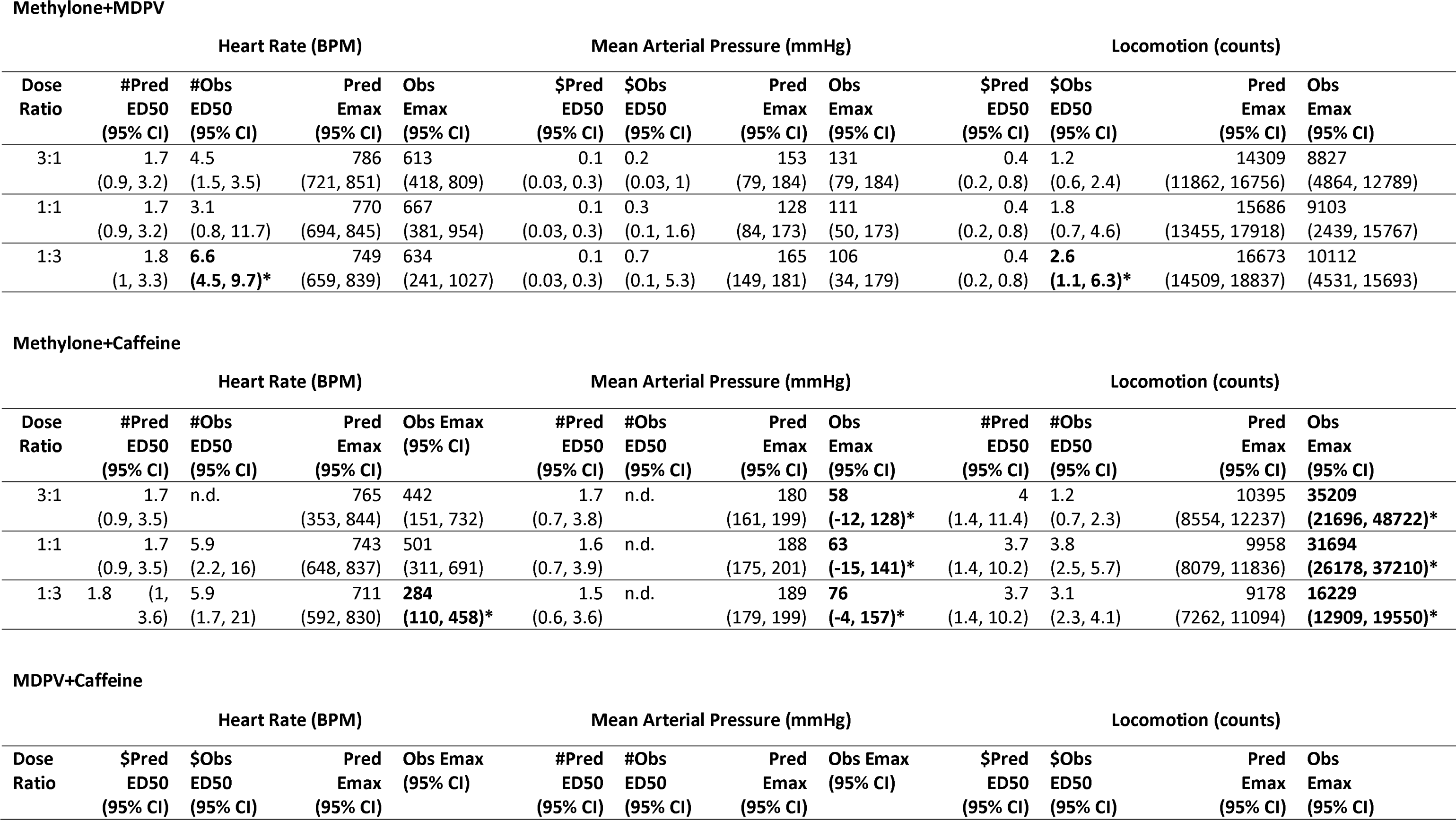
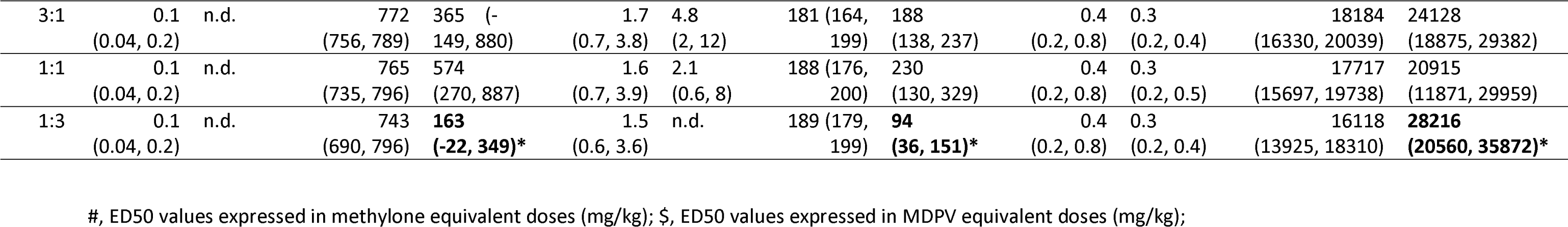
Predicted vs. observed potency and effectiveness values expressed as mean ± 95% confidence intervals. Bold values marked with an asterisk represent non-overlapping confidence intervals.

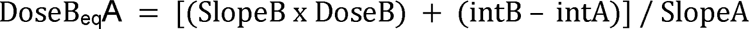

where SlopeA and SlopeB are the slope parameters and intA and intB are the y-intercepts for the drug producing the greater effect and the drug producing the lesser effect for a given endpoint, respectively, and derived from the dose-effect curves of each drug (AUC for each dose of each drug normalized to the AUC of saline). The sum of DoseA and DoseB_eq_A (Dose_eq_A) represents the total equivalent dose for a dose-pair, expressed in terms of the drug with a greater maximal effect for a given endpoint. The contribution of the least effective drug to the predicted effect level was capped at DoseB_eq_A that produces its maximum effect level (e.g., if Drug B produces 75% of the maximal effect level of Drug A, its contribution to the predicted effect level was capped at the dose of Drug A that produces 75% of the maximal effect). Dose_eq_A is then used to calculate the predicted effect level for an additive interaction using the following equation:

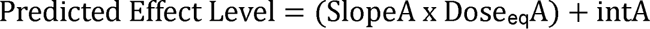

To evaluate the nature of the interaction for binary mixtures, experimentally determined ED_50_ (95% CIs) and E_max_ values for each binary mixture were compared to the ED_50_ (95% CIs) or E_max_ (95% CIs) derived from the dose-effect curve predicted for an additive interaction (i.e., predicted versus observed ED_50_ and E_max_ values). When 95% CIs overlapped, the interaction was considered to be strictly additive; however, when 95% CIs did not overlap the interaction was considered to be either supra-additive (mean ED_50_ was smaller or E_max_ was larger than predicted) or sub-additive (mean ED_50_ was larger or E_max_ was smaller than predicted). Prism 9 software (GraphPad Software Inc., La Jolla, CA, USA) was used to conduct statistical analyses and plot figures.

### Pharmacokinetics

To evaluate the pharmacokinetic profiles of methylone, MDPV, and caffeine, rats were treated with one dose of either MDPV (0.032-3.2 mg/kg; n=3), methylone (0.1-10 mg/kg; n=3), or caffeine (0.32-32 mg/kg; n=2) via infusion through one of the two catheters. Blood (0.4 ml) was then drawn through the alternate catheter 5-min, 60-min, 120-min, and 240-min following drug infusion; blood volumes were replaced with sterile saline. Different groups of rats were used for each drug, with the doses of each drug evaluated in a pseudorandom order. Subsequently, a separate group of rats was administered the largest dose-pairs of methylone+MDPV (n=3) and methylone+caffeine (n=3) evaluated in telemetry studies in order to detect pharmacokinetic interactions between these drugs. For those evaluations, blood (0.4 ml) was collected 5-min, 60-min, 120-min, and 240-min following drug infusion; blood volumes were replaced with sterile saline.

Pharmacokinetic data are presented as mean ± S.E.M. and dose-effect curves were generated using the area under the curve for each dose or drug mixture. The predicted area under the curve values for each dose were generated by multiplying the area of the curve produced by the smallest dose of each drug by the ratio of the dose of interest to the smallest dose evaluated (i.e., to calculate the predicted area under the curve for 1 mg/kg, multiply the area under the curve generated by 0.1 mg/kg by 10), as described previously (Anizan et al., 2016). Predicted versus observed area under the curve values were analyzed via unpaired t-tests.

#### HPLC/MS/MS System, Analytical Parameters, and Sample Extraction

The high-pressure liquid chromatography (HPLC) system consisted of a Shimadzu DGU-20A3R degassing unit (Shimadzu Scientific Instruments, Columbia, MD, USA), two LC-20AD pumps, a SIL-20A/HT autosampler, a CTO-20A oven (maintained at 26°C), and an AB Sciex API 3200 tandem mass spectrometer with turbo ion spray (AB Sciex, Framingham, MA, USA). The analytical column was an Ace Excel 3 C18-PFP (3L×L75Lmm, 3 µm) purchased from MacMod (Chadds Ford, PA, USA).

Mobile phase A was 0.1% formic acid (Sigma, St. Louis, MO) and 99.9% Milli-Q water (Millipore, Billerica, MA). Mobile phase B was 0.1% formic acid and 99.9% LCMS-grade acetonitrile (Thermo Fisher, Waltham, MA). The flow rate of the mobile phase was 0.4Lml/min. The gradient used was 0–1 min, 0% B; 1.1–5 min, 0–100% B (linear); 5.1–12 min, 100% B; 12.1–15 min, 0% B. The assay was performed in positive MRM mode. The Q1/Q3 transitions used to quantify methylone, caffeine, and MDVP were 208.0–160.0 m/z, 195.0–138.0 m/z, and 276.0–126.0 m/z, respectively. Internal standards D3-methylone, D3-caffeine, and D8-MDVP were used to control for variations in extraction efficiency with Q1/Q3 transitions of 211.0–135.1 m/z, 198.3–42.2 m/z, and 284.2–134.2 m/z, respectively.

Methylone, caffeine, and MDVP super-stock solutions were prepared in methanol at a concentration of 1 mg/ml and stored in aliquots at −80 °C. Deuterated methylone, caffeine, and MDVP super-stock solutions were prepared in methanol at concentrations of 100 µg/ml, 1 mg/ml, and 100 µg/ml, respectively, and stored in aliquots at −80 °C. A working stock solution mix containing methylone, caffeine, and MDVP (1, 10, and 100 µg/ml in methanol) was prepared fresh the day of the assay from the super-stock solutions and was used to spike the calibrator samples. A working stock solution mix of the internal standards was also prepared fresh on the day of the assay at the following concentrations (in methanol): D3-methylone, 8.33 µg/ml; D3-caffeine, 133.33 µg/ml; and D8-MDVP, 8.33 µg/ml. 15 µl of internal standard solution mix was added to all calibrators and unknown whole blood samples and vortexed for 15 seconds. To determine whole blood concentrations of methylone, caffeine, and MDVP, 200 µl of the calibrator and unknown whole blood samples were crashed with 640 µl of 0.1 % formic acid: 99.9% ethanol, vortexed for 15 seconds, then additionally mixed with 160 µl of 0.1% formic acid: 99.9% Milli-Q water, and vortexed again for 15 seconds. The samples were shaken for 20 minutes on a platform shaker and then centrifuged at 3200G for 25 minutes at 5°C. Supernatants were transferred to new tubes and dried to residue in a 60°C water bath under a stream of nitrogen. Residues were resuspended in 100 µl of mobile phase A:mobile phase B (1:3), vortexed, and then centrifuged at 13,000G for 5 minutes at 22°C. 10 µl of the final samples were injected into the liquid chromatography–tandem mass spectrometry system.

For unknown samples, the peak area ratios of methylone/D3-methylone were compared against a linear regression of peak area ratios of the calibrator samples with an internal standard at concentrations of 0, 10, 50, 200, 600, 1000, and 5000 ng/ml to calculate methylone concentrations. The peak area ratios of caffeine/D3-caffeine were compared against a quadratic regression of peak area ratios of the calibrator samples with an internal standard at concentrations of 0, 10, 50, 200, 600, 1000, 5000, 10000, and 20000 ng/ml to calculate caffeine concentrations in unknown samples. The peak area ratios of MDVP/D8-MDVP were compared against a linear regression of peak area ratios calibrator samples with internal standards at concentrations of 0, 10, 50, 200, 600, and 1000 ng/ml to calculate MDVP concentrations in unknown samples.

## RESULTS

### Cardiovascular and Locomotor Effects of Methylone

**Error! Reference source not found** (upper left) shows methylone-induced changes in heart rate as a function of time, with a mixed-effects 2-way ANOVA revealing significant main effects of dose (F_(2.7, 21.5)_=12.2; p<0.0001) and time (F_(2.9, 23.4)_=18.8; p<0.0001). Consistent with this, a 1-way ANOVA revealed a significant effect of dose (F_(2.7, 120.7)_=12.6; p<0.0001) on the area under the curve for each dose of methylone (Figure 1; upper right). Whereas methylone produced significant increases in heart rate, a 2-way ANOVA of the time course (Figure 1; middle left) and a 1-way ANOVA of the area under the curve (Figure 1; middle right) failed to identify a significant effect of dose on mean arterial pressure (F_(2.4, 19.6)_=0.1; p>0.05) and (F_(2.4, 18.8)_=0.1; p>0.05), respectively. Figure 1 (lower panels) depicts the time course and area under the curve of locomotor activity elicited by methylone with a 2-way ANOVA revealing significant main effects of dose (F_(1.7, 14)_=11.3; p<0.01) and time (F_(1.7, 14)_=26; p<0.0001), and a 1-way ANOVA of the area under the curve revealing a significant effect of dose (F_(1.7, 13.3)_=23.7; p<0.0001).

**Figure 1:**
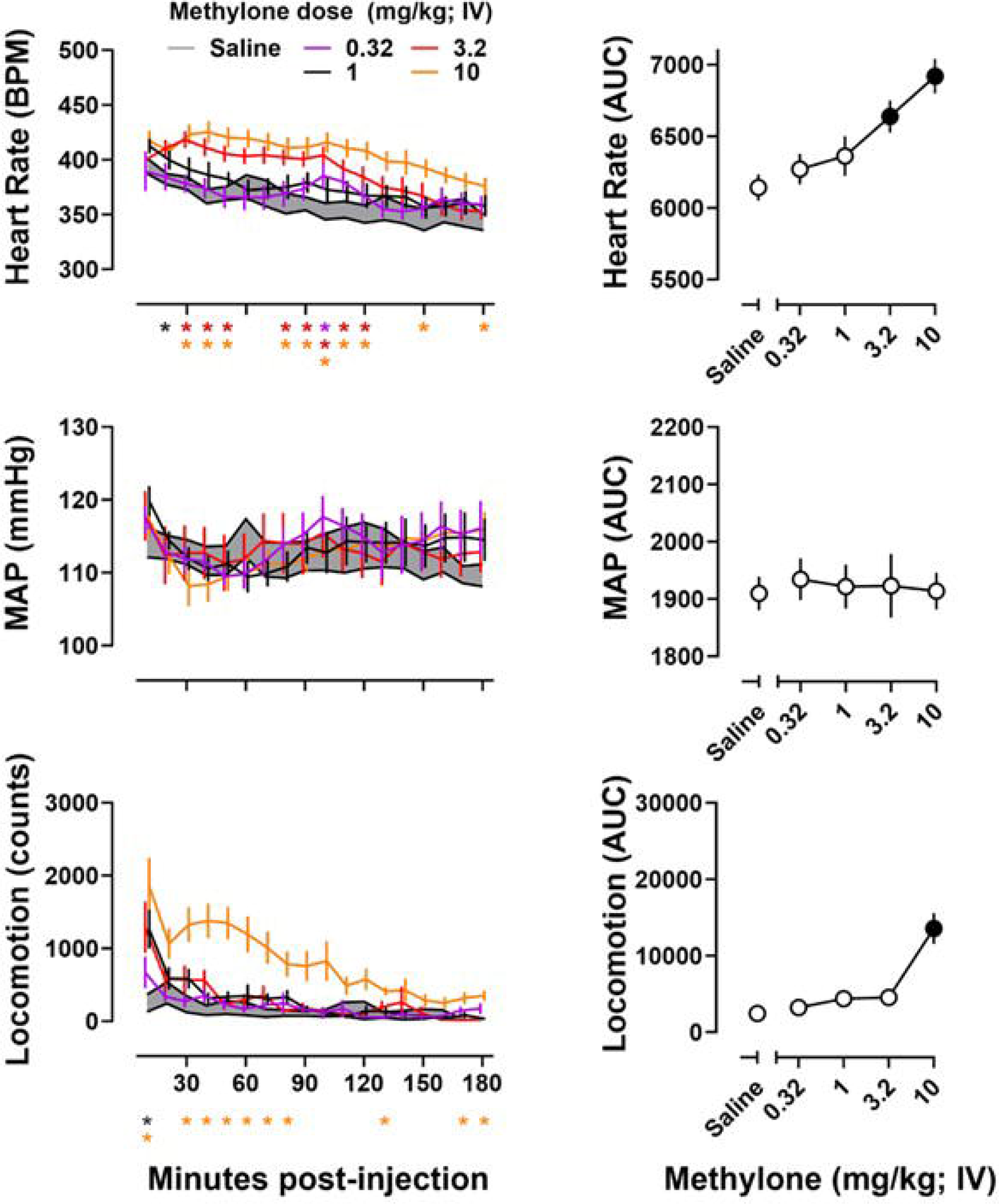
Time courses of the effects of increasing doses of methylone (mg/kg) on heart rate (upper left), mean arterial pressure (middle left), and locomotion (lower left). Asterisks represent a significant difference from saline at that time point corresponding with the dose associated with that color. * denotes p<0.05. Area under the curve of heart rate (top right), mean arterial pressure (middle right), and locomotion (bottom right) were calculated from the corresponding time course data. Data represent the mean ±S.E.M. Filled symbols represent a significant difference from saline.

### Cardiovascular and Locomotor Effects of MDPV

The cardiovascular and locomotor effects produced by MDPV are depicted in Figure 2. Figure 2 (upper left) shows the changes in heart rate produced by MDPV as a function of time, with a mixed-effects 2-way ANOVA revealing significant main effects of dose (F_(3.2, 29)_=5.3; p<0.01) and time (F_(1.9, 17.4)_=6.4; p<0.01). Consistent with this, the area under the curve calculations for each dose of MDPV (Figure 2; upper right) illustrate the dose-dependent increases produced by MDPV, with a 1-way ANOVA revealing a significant effect of dose (F_(3.3,29.5)_=5.4; p<0.01). Figure 2 (middle) depicts the effects on mean arterial pressure produced by MDPV. A 2-way ANOVA revealed significant effects of dose (F_(3.4, 30.5)_=4.5; p<0.01) and time (F_(2.1, 19.1)_=3.6; p<0.05) on mean arterial pressure. A 1-way ANOVA of the area under the curve data also revealed significant effects of dose (F_(3.3, 28.1)_=4.5; p<0.01). Figure 2 (lower panels) depicts the time course and area under the curve illustrating dose-dependent increases in locomotor activity produced by MDPV with a 2-way ANOVA revealing significant main effects of dose (F_(1.8,15.9)_=22.2; p<0.0001) and time (F_(3.8, 34.5)_=68.5; p<0.0001), and a 1-way ANOVA of the area under the curve for locomotor activity revealing a significant effect of dose (F_(1.8, 15.9)_=22.2; p<0.0001).

**Figure 2:**
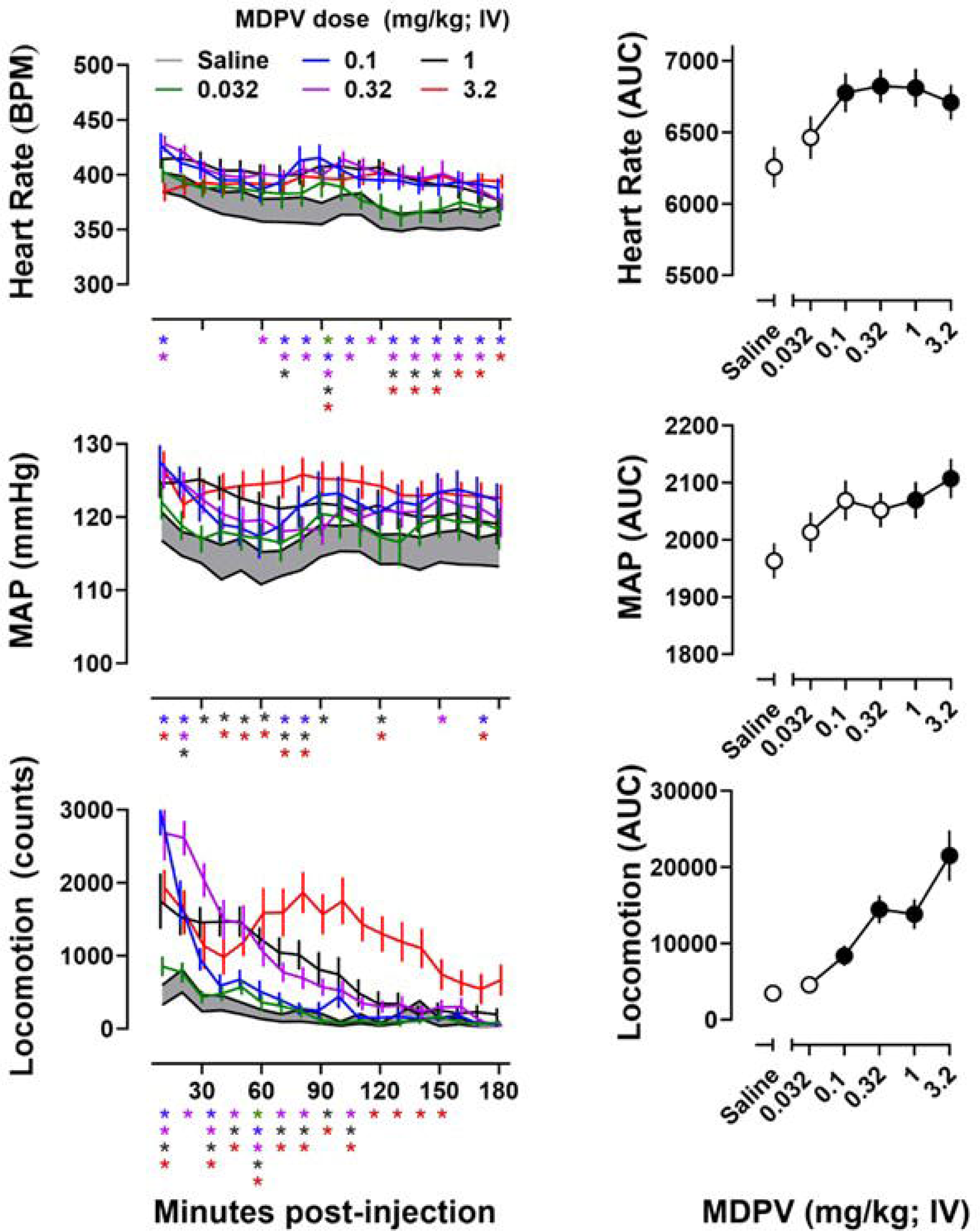
Time courses of the effects of increasing doses of MDPV (mg/kg) on heart rate (upper left), mean arterial pressure (middle left), and locomotion (lower left). Asterisks represent a significant difference from saline at that time point corresponding with the dose associated with that color. * denotes p<0.05. The area under the curve of heart rate (top right), mean arterial pressure (middle right), and locomotion (bottom right) were calculated from the corresponding time course data. Data represent the mean ±S.E.M. Filled symbols represent a significant difference from saline.

### Cardiovascular and Locomotor Effects of Caffeine

Figure 3 (upper panels) shows the effects on heart rate produced by caffeine as a function of time, although mixed-effects 2-way ANOVA of the time course of caffeine revealed no significant main effects of dose (F_(2.5,24.6)_=1.1; p>0.05) but a significant effect of time (F_(2.2,22.1)_=9.9; p<0.001). The area under the curve calculations for each dose of caffeine (Figure 3; upper right) illustrates the increases produced by caffeine, however, a 1-way ANOVA revealed no significant effect of dose (F_(2.5, 23.9)_=1; p=0.4). Similarly, Figure 3 (middle) depicts the dose-dependent increases in mean arterial pressure produced by caffeine as a function of time, however, mixed-effects 2-way ANOVA of the timecourse of caffeine revealed no significant main effects of dose (F_(2.1,20.9)_=3.1; p=0.06) nor time (F_(2.2,22.3)_=2.6; p>0.05). The area under the curve calculations for each dose of caffeine (Figure 4; middle right) illustrates the dose-dependent increases produced by caffeine on mean arterial pressure, but a 1-way ANOVA revealing only a trend towards a significant effect of dose (F_(2.1, 20.2)_=3.2; p=0.06). Locomotor effects of caffeine (Figure 3; lower panels) depict the time course and area under the curve of locomotor activity elicited by caffeine with a 2-way ANOVA revealing significant main effects of dose (F_(2.6, 26.4)_=14.2; p<0.0001) and time (F_(3.1,31.1)_=33.4; p<0.0001), and a 1-way ANOVA revealed a significant effect of dose (F_(2.6, 25.5)_=14; p<0.0001).

**Figure 3:**
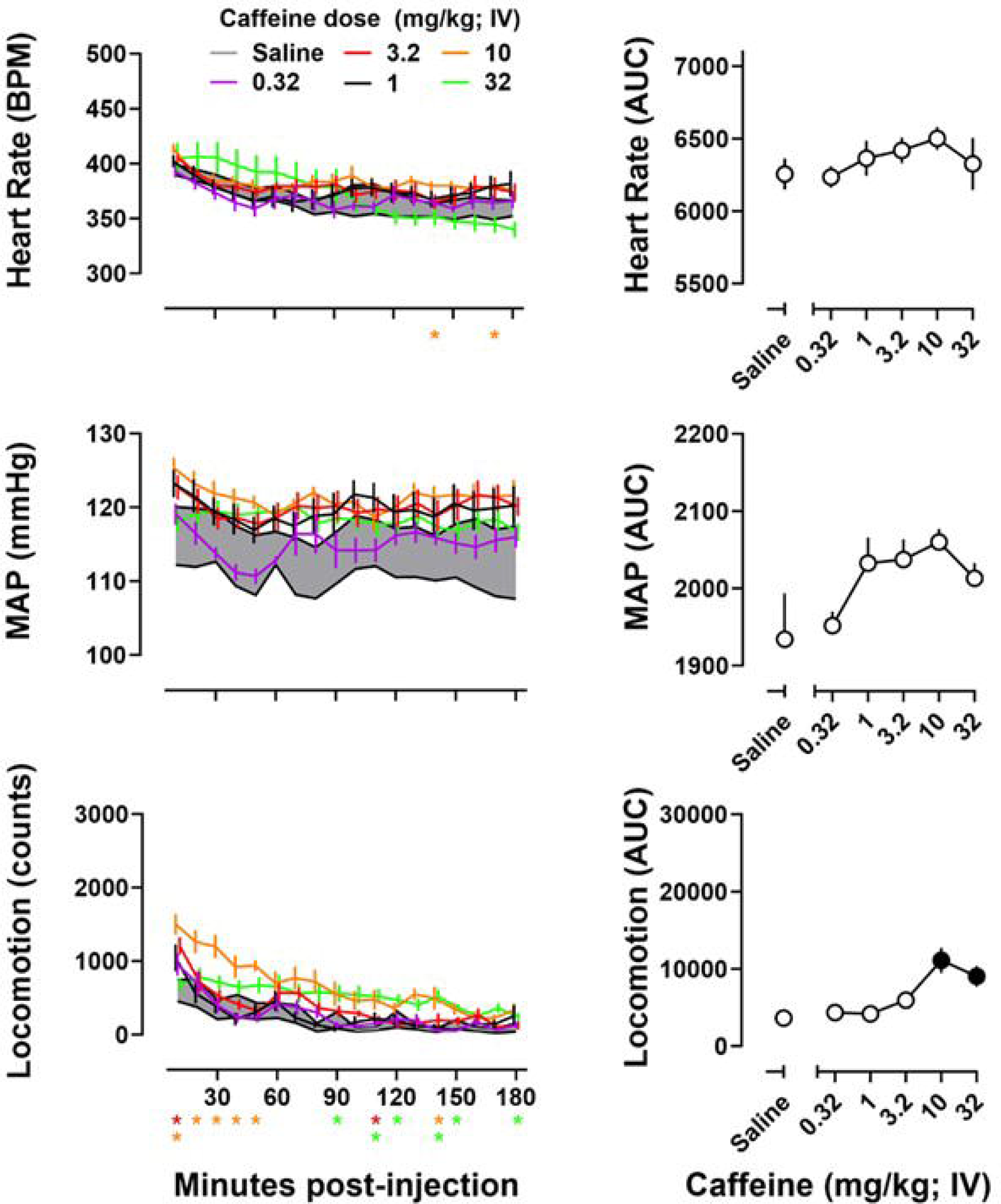
Time courses of the effects of increasing doses of caffeine (mg/kg) on heart rate (upper left), mean arterial pressure (middle left), and locomotion (lower left). Asterisks represent a significant difference from saline at that time point corresponding with the dose associated with that color. * denotes p<0.05. The area under the curve of heart rate (top right), mean arterial pressure (middle right), and locomotion (bottom right) were calculated from the corresponding time course data. Data represent the mean ±S.E.M. Filled symbols represent a significant difference from saline.

**Figure 4:**
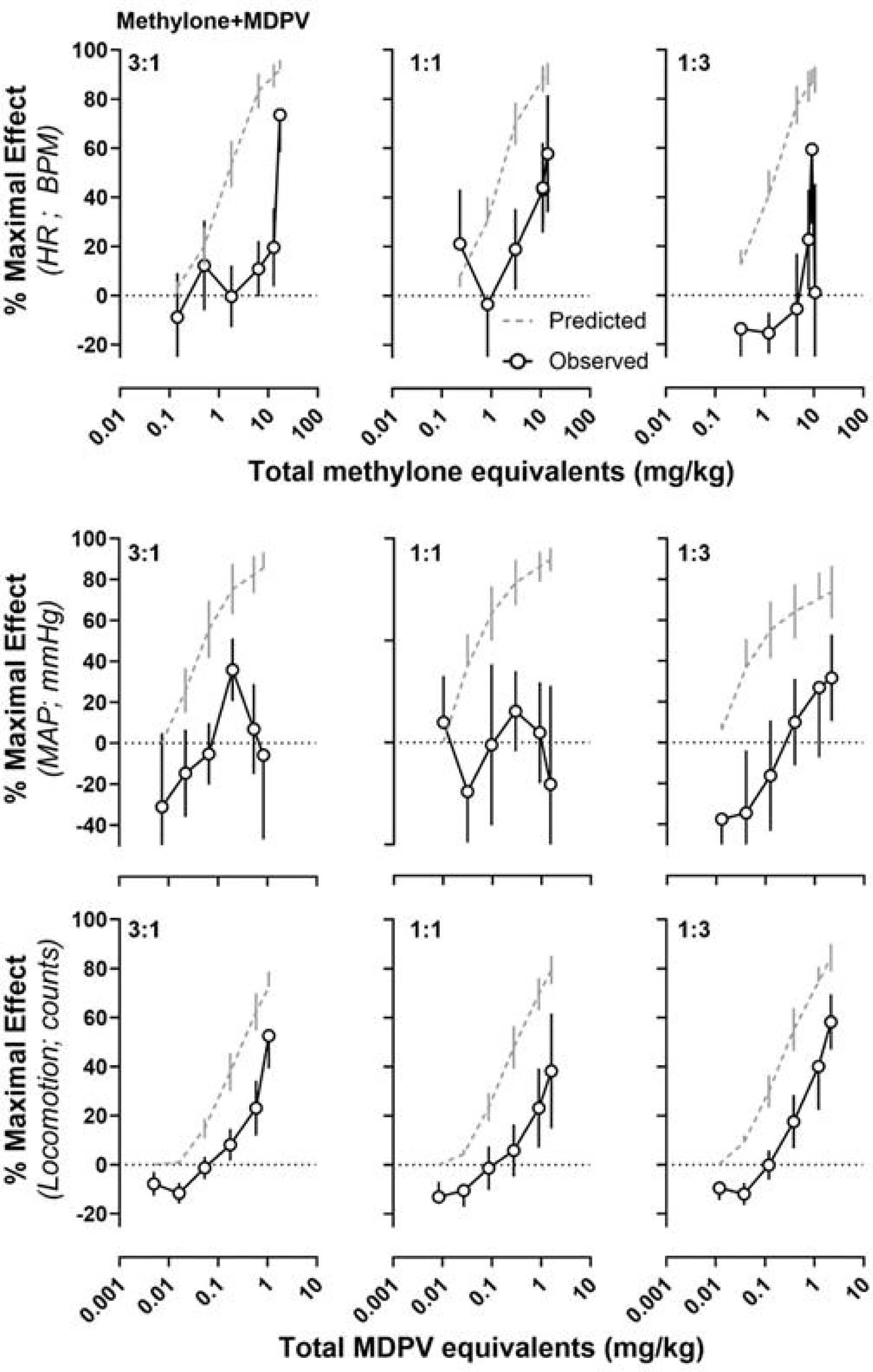
Experimentally determined (open circles) versus predicted, additive (grey, dashed lines) dose-effect curves for the effects of binary mixtures of methylone+MDPV on heart rate (upper panels), mean arterial pressure (middle panels), and locomotion (lower panels). The ordinate represents the percent of the maximal effect. Data represent the mean ±S.E.M.

### Cardiovascular and Locomotor Effects of Binary Mixtures of Methylone and MDPV

Predicted and observed dose-effect curves for mixtures of methylone+MDPV are shown in Figure 4. The mean ED_50_ and E_max_ values (±95% confidence intervals) for each mixture of methylone+MDPV are reported in Table 2 (top row). Overall, mixtures of methylone+MDPV were less potent and effective than predicted, resulting in significant sub-additive interactions with regard to potency to increase heart rate for the 1:3 mixture, and with regard to the maximal effect of the 1:1 mixture on mean arterial pressure. Administration of the largest dose pair of the 3:1 methylone+MDPV mixture resulted in convulsions in 2/9 rats and in one of those cases, lethality.

### Cardiovascular and Locomotor Effects of Binary Mixtures of MDPV and Caffeine

Predicted and observed dose–effect curves for mixtures of MDPV+caffeine are shown in Figure 5. The mean ED_50_ and E_max_ values (±95% confidence intervals) for each mixture of MDPV+caffeine are reported in Table 2 (middle row). Overall, mixtures of MDPV+caffeine did not depart from additivity with regard to potency to increase heart rate, but the 1:3 mixture did result in a significant sub-additive interaction with regard to maximal effect. With regard to locomotor activity, MDPV and caffeine exhibited largely additive interactions with respect to their potency but increases in maximal effect were observed, with the 1:3 mixture resulting in a significant supra-additive interaction. Administration of the largest dose pair of MDPV+caffeine produced convulsions in 1/8 rats following administration of the 3:1 mixture and 2/9 rats following administration of the 1:3 mixture.

**Figure 5:**
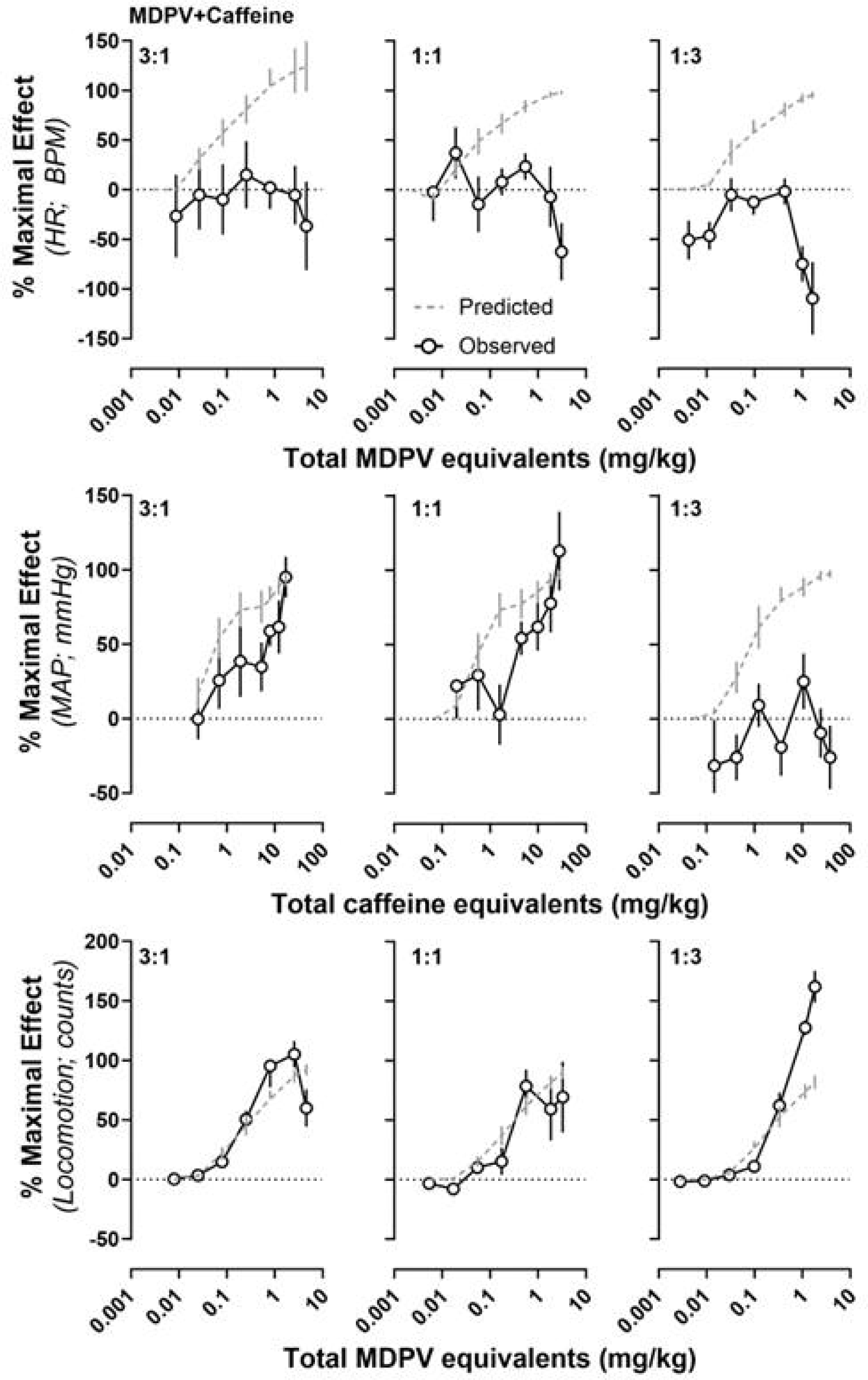
Experimentally determined (open circles) versus predicted, additive (grey, dashed lines) dose-effect curves for the effects of binary mixtures of MDPV+caffeine on heart rate (upper panels), mean arterial pressure (middle panels), and locomotion (lower panels). The ordinate represents the percent of the maximal effect. Data represent the mean ±S.E.M.

### Cardiovascular and Locomotor Effects of Binary Mixtures of Methylone and Caffeine

Predicted and observed dose–effect curves for mixtures of methylone+caffeine are shown in Figure 6. The mean ED_50_ and E_max_ values (±95% confidence intervals) for each mixture of methylone+caffeine are reported in Table 2 (bottom row). With regard to effects on heart rate and blood pressure, mixtures of methylone+caffeine resulted in sub-additive interactions with regard to potency and effectiveness. In contrast, mixtures of methylone+caffeine resulted in significant supra-additive interactions with regard to increases in locomotor activity (without significantly changing potency). Administration of the largest dose pair of methylone+caffeine mixtures produced convulsions in 4/8 rats following the 3:1 mixture, 3/9 rats following the 1:1 mixture, and 4/10 rats (with two deaths occurring) following the 1:3 mixture.

**Figure 6:**
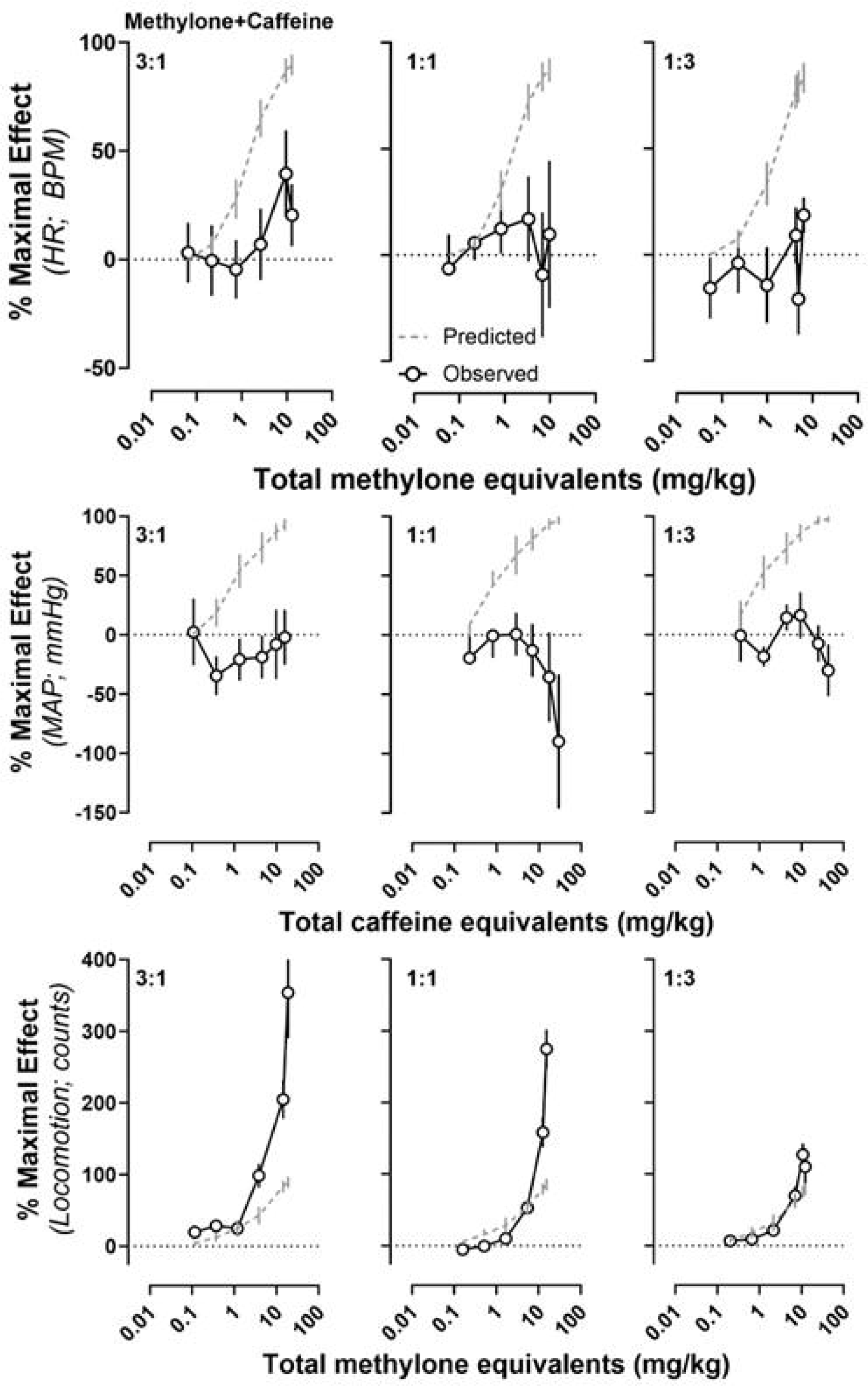
Experimentally determined (open circles) versus predicted, additive (grey, dashed lines) dose-effect curves for the effects of binary mixtures of methylone+caffeine on heart rate (upper panels), mean arterial pressure (middle panels), and locomotion (lower panels). The ordinate represents the percent of the maximal effect. Data represent the mean ±S.E.M.

#### Pharmacokinetics

Figure 7 depicts the time courses of blood levels of methylone (upper left) and the predicted versus observed area under the curve values for methylone (upper right), with the largest dose of methylone (10 mg/kg) resulting in a significantly greater area under the curve value than the predicted area under the curve value (p<0.05). Also depicted are the time courses of blood levels and the predicted versus observed area under the curve values for MDPV (Figure 7; middle panels) and caffeine (Figure 7; lower panels). The area under the curve produced by the largest dose of caffeine (32 mg/kg) was significantly smaller than the predicted area under the curve calculated for that dose of caffeine (p<0.05). Figure 8 (left) illustrates the predicted and observed area under the curve values for methylone and MDPV when administered in the largest 3:1 dose pair of the methylone+MDPV mixture that was evaluated in telemetry. Consistent with what was observed when the drugs were administered alone, the observed area under the curve value for methylone was significantly greater than the predicted value (p<0.05), whereas no significant difference between the predicted and observed area under the curve values was observed for MDPV. Similarly, Figure 8 (right) shows the predicted and observed area under the curve values for methylone and caffeine when administered in the largest 3:1 dose pair of the methylone+caffeine mixture that was evaluated in telemetry. Consistent with what was observed when the drugs were administered alone, the observed area under the curve value for methylone was significantly greater than the predicted value (p<0.05), whereas no significant difference between the predicted and observed area under the curve values was observed for caffeine.

**Figure 7:**
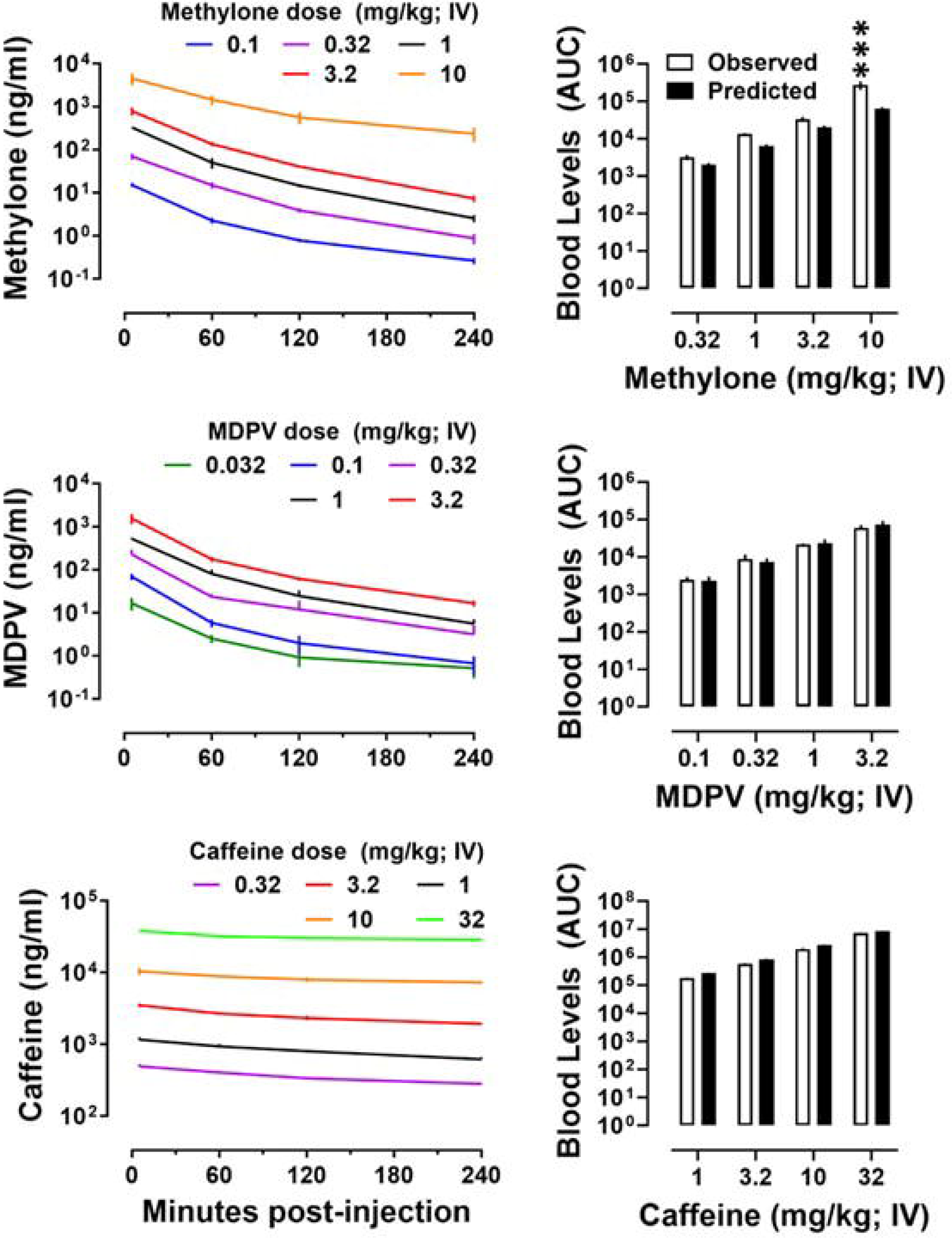
Blood levels of increasing doses (expressed as mg/kg) of methylone (upper left), MDPV (middle left), and caffeine (lower left) as a function of time and experimentally determined area under the curve (right; open bars) versus predicted area under the curve (right; filled bars). Data represent the mean ±S.E.M. ** signifies p<0.01. *** signifies p<0.001.

**Figure 8:**
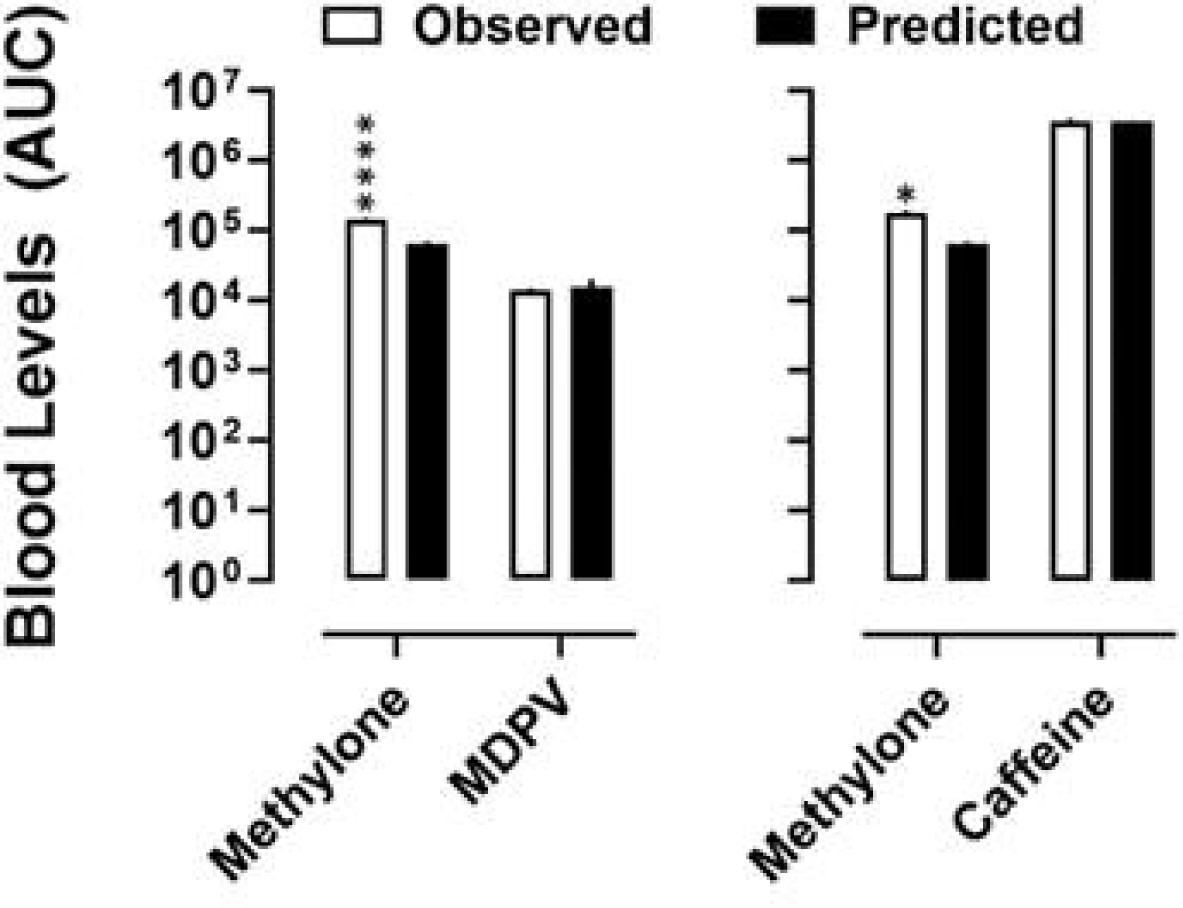
Experimentally determined area under the curve (open bars) versus predicted area under the curve (filled bars) for methylone and MDPV when co-administered (left) and for methylone and caffeine when co-administered (right). Data represent the mean ±S.E.M. * denotes p<0.05. **** denotes p<0.0001.

## DISCUSSION

Although “Bath Salts” preparations commonly contain mixtures of multiple synthetic cathinones, or synthetic cathinones and caffeine, little is known about the manner in which constituents interact to produce adverse physiological and psychological effects. To this end, the current studies evaluated the cardiovascular and locomotor effects of three common “Bath Salts” constituents, methylone, MDPV, and caffeine, and utilized dose-addition analyses to define the nature of the interactions between binary “Bath Salts” mixtures comprising two synthetic cathinones (i.e., methylone+MDPV) and either synthetic cathinone and caffeine (i.e., methylone+caffeine and MDPV+caffeine). There were 5 main findings: 1) methylone produced increases in heart rate and locomotor activity whereas MDPV and caffeine produced increases in heart rate, as well as mean arterial pressure, and locomotor activity; 2) mixtures of methylone+MDPV resulted in sub-additive to additive interactions with regard to heart rate, blood pressure, and locomotor activity whereas mixtures of either cathinone+caffeine also resulted in sub-additive to additive interactions with regard to heart rate and mean arterial pressure but additive to supra-additive interactions with regard to locomotor activity; 3) although convulsions and lethality were not observed following administration of any drugs alone, some convulsions occurred following administration of the largest dose pairs of “Bath Salts” mixtures (methylone+caffeine > MDPV+caffeine > methylone+MDPV), with lethality observed following administration of the largest dose pair of the 1:3 mixture of methylone+caffeine; 4) methylone, MDPV, and caffeine exhibited largely linear pharmacokinetics, with the exception that the largest dose of methylone resulted in greater than predicted blood levels of methylone; and 5) the largest dose pairs of 3:1 methylone+MDPV and 3:1 methylone+caffeine did not alter the blood levels of either constituent relative to predicted blood levels. Together, these data provide direct evidence that the constituents present in a given “Bath Salts” preparation can significantly impact cardiovascular parameters (i.e., heart rate, mean arterial pressure), locomotor activity, and the likelihood of lethality.

Consistent with previous studies in rats, MDPV and caffeine produced dose-related increases in heart rate, mean arterial pressure, and locomotion (Ilbäck et al., 2007; Pechlivanova et al., 2010; Han et al., 2011; Baumann et al., 2013; Schindler et al., 2016b; McClenahan et al., 2019). Regarding methylone, the dose-dependent increases in locomotor activity observed in the current studies are consistent with other work demonstrating the locomotor stimulating effects of methylone (Elmore et al., 2017; Javadi-Paydar et al., 2018; Centazzo et al., 2021); however, to our knowledge, this is the first preclinical study to evaluate the cardiovascular effects of methylone, wherein we found that methylone produced a marked increase in heart rate without significant effects on mean arterial pressure. The rank order for increases in heart rate was methylone > MDPV > caffeine, whereas the rank order for increases in mean arterial pressure was caffeine > MDPV > methylone, and the rank order for increases in locomotor activity was MDPV > methylone > caffeine.

Consistent with sub-additive interactions with regard to reinforcing effects (Gannon et al., 2018b), mixtures of methylone and MDPV were largely sub-additive with regard to potency and effectiveness for producing effects on heart rate, mean arterial pressure, and locomotor activity. In the current studies relying on bolus, non-contingent infusions, the largest dose pair of methylone+MDPV produced convulsions in 2 rats, one of which subsequently died. Importantly, this is also consistent with the results of our previous study (Gannon et al., 2018b), in which lethality was observed in approximately 30% of rats that self-administered mixtures that contained large doses of methylone and MDPV. Together, these observations suggest that the sub-additive interactions observed between the reinforcing and cardiovascular effects of methylone and MDPV are unlikely to have resulted from large reductions in the potency of the mixture to produce any effect. Importantly, it is well known that the nature of the interaction between two drugs can vary across endpoints, and although cardiovascular and locomotor endpoints were selected as proxies for the types of adverse effects reported in humans, it seems likely that interactions between other effects of methylone and MDPV are driving the observed increases in overt toxicity. To investigate whether pharmacokinetic interactions could have contributed to the effects produced by methylone+MDPV mixtures, blood levels of methylone and MDPV were quantified after administration of the largest dose pair of the 3:1 ratio of methylone+MDPV and compared the blood levels predicted based on the pharmacokinetic profiles of each drug alone. Similar to what we observed following the administration of a 10 mg/kg dose of methylone alone, blood levels of methylone were significantly greater than predicted for linear pharmacokinetics when the largest dose of methylone was administered in combination with MDPV. This is consistent with previous reports that methylone acts as a mechanism-based inactivator of CYP2D6, the enzyme primarily responsible for its metabolism (Pedersen et al., 2013; Elmore et al., 2017). In contrast, blood levels of MDPV did not differ from predicted levels when administered alone, or in combination with methylone, suggesting that observed interactions between MDPV and methylone were not pharmacokinetic in nature. Since MDPV has been shown to be metabolized by CYP1A2 and CYP2C19, in addition to CYP2D6, it is likely that the inactivation of CYP2D6 by methylone was insufficient to alter its linear pharmacokinetic profile (Meyer et al., 2010; Strano-Rossi et al., 2010; Anizan et al., 2016) at the doses evaluated. Importantly, though the current study used blood levels of MDPV and methylone to investigate possible pharmacokinetic interactions, future studies should investigate the potential for pharmacokinetic interactions in brain tissue, as interactions here might be more tightly linked to observed changes in behavioral and physiologic effects of MDPV and methylone.

Although administration of either drug, in isolation, produced comparable effects on heart rate and locomotor activity, MDPV acting as a monoamine reuptake inhibitor could have interfered with the effects of methylone, which acts as a substrate for monoamine transporters, and vice-versa, resulting in smaller effects than predicted for the doses evaluated in the current study. Although this proposed mechanism underlying the sub-additivity observed for cardiovascular and locomotor effects is speculative, others have reported that MDPV can inhibit methamphetamine-induced dopamine efflux and prevent the neurotoxic effects of methamphetamine (Simmler et al., 2013; Anneken et al., 2015). Perhaps MDPV interfering with the catecholaminergic effects of methylone was sufficient to result in reduced effects across each parameter evaluated in the current studies that are largely reliant upon catecholaminergic activity.

Given that supra-additive interactions have been observed between cathinones and caffeine with regard to reinforcing and discriminative stimulus effects (Collins et al., 2016; Gannon et al., 2018b, 2019; Doyle et al., 2021), it was unexpected that the interactions between the cardiovascular effects of the synthetic cathinones and caffeine were largely sub-additive. Mixtures of methylone and caffeine resulted in sub-additive interactions for both heart rate and blood pressure, whereas mixtures of MDPV and caffeine resulted in sub-additive interactions for heart rate, but additive interactions for blood pressure (with the exception of 1:3 MDPV+caffeine, which resulted in a sub-additive interaction with regard to blood pressure). In contrast to the cardiovascular effects, additive to supra-additive interactions were observed between the locomotor effects of either synthetic cathinone and caffeine. These findings are consistent with previous work from our laboratory demonstrating supra-additive interactions between methylone and caffeine, and MDPV and caffeine, with regard to reinforcing effectiveness using a progressive ratio schedule of reinforcement and demand curve analyses (Gannon et al., 2018b, 2019). Although a rigorous evaluation of the mechanisms underlying departures from additivity between either synthetic cathinone and caffeine was outside the scope of the current study, one possible explanation for the observation of sub-additive interactions for cardiovascular effects, but supra-additive interactions for locomotor effects is the relationship between dopamine D_2_ receptor signaling and that of adenosine A_2A_ receptors. Previous work has demonstrated that dopamine D_2_ and adenosine A_2A_ receptors form heteromeric complexes and that tonic activation of A_2A_ receptors by adenosine is capable of dampening signaling through dopamine D_2_ receptors (Fuxe and Ungerstedt, 1974; Ferre et al., 1991; Ferré et al., 1997, 1997; Ferré, 2016). Indeed, the disinhibition of dopamine D_2_ receptor signaling is thought to be the primary mechanism through which caffeine stimulates locomotor activity (Fuxe and Ungerstedt, 1974; Ferre et al., 1991; Ferré et al., 1997; Ferré, 2016). Through the antagonism of adenosine receptors, caffeine can increase dopamine D_2_ receptor signaling which has been suggested to dampen noradrenergic tone and result in sympatholytic effects (Cavero et al., 1982; Barrett and Lokhandwala, 1983; Goldberg and Murphy, 1987; Lefevre-Borg et al., 1987; Velasco and Luchsinger, 1998; Jose et al., 2003; Zeng et al., 2007). Thus, enhancements of dopamine D_2_ receptor signaling could be responsible for both the sub-additive interactions observed with regard to cardiovascular effects and supra-additive interactions observed with regard to locomotor and reinforcing effects. Of note, it is unlikely that pharmacokinetic interactions between either cathinone and caffeine are driving the observed effects, as we demonstrated no changes in the blood levels of caffeine when co-administered with a large dose of methylone. With regard to caffeine and MDPV, each is metabolized by a host of enzymes and has not been demonstrated to inactivate their own metabolisms, making it unlikely that a pharmacokinetic interaction underlies the observed departures from additivity between these two drugs (Kot and Daniel, 2008; Anizan et al., 2016). Strikingly, the largest dose pairs of methylone+caffeine mixtures resulted in 11 rats convulsing (4 following the 3:1 ratio, 3 following the 1:1 ratio, and 4 following the 1:3 ratio) with death occurring in 2 rats (both following administration of the largest dose pair of the 1:3 methylone+caffeine mixture). No convulsions and/or deaths were observed following the administration of any drug alone in the current studies, suggesting a potential synergism with regard to toxicity (i.e., convulsions) and/or lethality produced by methylone and caffeine. This interaction is likely not a result of a pharmacokinetic interaction, since there were no changes in the blood levels of methylone or caffeine when co-administered, suggesting a potential pharmacodynamic interaction underlying these adverse effects. These data are largely consistent with previous work from our laboratory that investigated the reinforcing effectiveness of methylone+caffeine and MDPV+caffeine mixtures in rats and found that response-contingent access to drug mixtures containing methylone resulted in a relatively large number of deaths (Gannon et al., 2018b, 2019). Given reports of methylone being associated with toxicity and lethality in human “Bath Salts” users, and the prevalence of caffeine in these mixtures (Pearson et al., 2012), deeper investigation of the lethal interactions between these constituents is warranted.

Toxicity associated with the use of “Bath Salts” gained much attention due to the wide variety of reported bizarre and adverse effects that are not typical of other drugs with similar mechanisms of action. While previous studies have identified supra-additive interactions between the abuse-related effects of “Bath Salts” constituents (Collins et al., 2016; Gannon et al., 2018b; Doyle et al., 2021), the degree to which these interactions extend to physiological effects is not well understood. The present study directly assessed the cardiovascular and locomotor effects of mixtures of common “Bath Salts” constituents and found that drug preparations containing methylone+MDPV were less effective at stimulating heart rate, blood pressure, and locomotion than predicted based on an additive interaction, but produced incidents of convulsions and lethality that were not observed with either drug alone whereas drug preparations containing caffeine resulted in sub-additive to additive interactions with regard to cardiovascular effects, but additive to supra-additive interactions with regard to locomotor effects. Together, these studies suggest that the composition of “Bath Salts” preparations can have a significant impact on their cardiovascular and locomotor effects and suggest that such interactions could contribute to the adverse and lethal effects reported in human users of these drug preparations.

## ACKNOWLEDGEMENTS

The authors would like to thank Riley Pritchett, Rachel DeSantis, Christina George and Kelly Salinas for their technical assistance in the completion of these studies.

